# ibSLS: A Biobank for Democratizing Access to Multi-Omics Data and Biospecimens from Spaceflight Research

**DOI:** 10.1101/2025.09.08.675003

**Authors:** Akihito Otsuki, Yuichi Aoki, Risa Okada, Daisuke Kamimura, Dai Shiba, Eiji Hishinuma, Seizo Koshiba, Fumiki Katsuoka, Kengo Kinoshita, Takafumi Suzuki, Akira Uruno, Masayuki Yamamoto

## Abstract

During spaceflight, astronauts are exposed to extreme conditions such as microgravity, cosmic radiation, and confinement, which can cause a wide range of health problems. To elucidate the molecular mechanisms underlying these issues and to develop intervention strategies for maintaining physiological homeostasis during space missions, space life science research using mouse models is actively conducted on the International Space Station (ISS). However, because of the high cost and technical complexity of spaceflight experiments, it is critical to maximize the scientific value of each mission by ensuring broad accessibility to both data and biospecimens. To this end, we developed the integrated biobank for Space Life Sciences (ibSLS; https://ibsls.megabank.tohoku.ac.jp), a data-visualization and sample-sharing platform that provides access to transcriptomic and metabolomic datasets generated from JAXA’s Mouse Habitat Unit (MHU) missions. The platform features a user-friendly interface, tools for cross-mission analysis, and integration with human multi-omics databases, enabling cross-species comparisons. In addition, ibSLS facilitates biospecimen requests to support downstream research. By promoting open access to spaceflight-derived data and biological resources, ibSLS encourages the participation of researchers from diverse fields in space life science. We believe that ibSLS will make a valuable contribution to both biomedical research on spaceflight-related health issues and the study of diseases on Earth.

## Introduction

Since the first successful crewed spaceflight conducted in 1961^1^, outer space has been the frontier for the expansion of human activity. In recent years, not only have traditional government programs continued to explore space, but private companies have also begun to play a significant role, with growing interest in commercial applications and private-sector-led missions such as Inspiration4^2^. As access to space becomes more diversified, opportunities for human spaceflight are rapidly expanding.

During spaceflight missions, astronauts are exposed to extreme environmental conditions, including microgravity and high-dose cosmic radiation, which significantly impact physiological homeostasis^3^. To ensure the safety of future human missions and to deepen our understanding of biological responses to space environments, it is crucial to investigate the molecular effects of spaceflight and to develop strategies for mitigating them. To this end, basic life science experiments become critical, which are actively conducted in low Earth orbit, particularly aboard the International Space Station (ISS)^2^.

Given the difficulty of obtaining human tissue samples and inherent limitations in studying human biology in space, a variety of model organisms have been used in spaceflight experiments^4^. Among these, mouse models are particularly effective for the understanding of biological processes relevant to human health. To conduct precise mouse experiments on the ISS, the Japan Aerospace Exploration Agency (JAXA) developed a specialized experimental platform named the Multiple Artificial Gravity Research System (MARS)^5, 6, 7, 8^. MARS consists of several key components, including the Transfer Cage Unit (TCU), the Habitat Cage Units (HCU), and the Centrifuge-equipped Biological Experiment Facility (CBEF). The TCU is used to transport mice between Earth and the ISS, while the HCU is used to house them on board. Both systems ensure mouse housing in separate cages, thereby supporting controlled experimental conditions. By mounting the HCU onto the CBEF, artificial gravity can be applied via centrifugal force, allowing direct comparisons between microgravity and artificial gravity conditions on the ISS^5, 6^.

Using MARS, JAXA has conducted a series of missions known as the Mouse Habitat Unit (MHU) missions. These missions have successfully characterized molecular alterations in mouse tissues and cells that occur during and after spaceflight. A notable finding is the marked acceleration of aging-like phenotypes such as reductions in bone density and muscle mass during spaceflight, which were mitigated by applying artificial gravity^5, 9, 10^. Comprehensive transcriptomic and metabolomic analyses further revealed that spaceflight induces aging-like changes in plasma metabolites^11^. These insights highlight the value of rodent models in studying spaceflight-associated and age-related biological changes, providing critical knowledge for human spaceflight safety and for understanding aging-related disease mechanisms on Earth.

However, such space life science research often encounters substantial limitations compared with typical life science experiments on the ground. These include the scale and chance of experiments that can be carried out, the size and accessibility of experimental equipment, and the availability of trained staff for the experiment. Additionally, there are significant technical challenges in the development of experimental hardware suited to the extreme conditions os space, as well as in the optimization of sample collection, processing, and storage. Consequently, the opportunities for conducting life science experiments in space are limited. It is therefore of the utmost importance to maximize the scientific value of data and specimens recovered from each spaceflight experiment.

To organize the results of the costly space experiments conducted in the MHU missions and to make the omics data and biospecimens widely available, we developed the integrated biobank for Space Life Science (ibSLS; https://ibsls.megabank.tohoku.ac.jp). Since the launch of ibSLS in 2020, the biobank has provided researchers with a variety of omics data from the MHU missions, along with related metadata and accession to the biological sample-sharing program conducted by JAXA. Furthermore, by linking with a large human cohort database developed by the Tohoku Medical Megabank Project^12, 13, 14, 15, 16, 17^ in Japan, ibSLS enables researchers to extrapolate findings from MHU missions to interpretation of human phenotypes. Here, we describe the concepts, architecture, and contents of the ibSLS, as well as its ongoing efforts in integrated analysis and data visualization to support future space biology research.

## Results

### Concept of the integrated biobank for space life science (ibSLS)

The space environment exerts a considerable influence on multiple biological systems, including bones, muscles, heart, and blood flow, in astronauts and space travellars^3^. To elucidate molecular mechanisms underlying these biological changes, experimental approaches utilizing mouse models are particularly valuable. Compared to other model organisms, such as nematodes or plants that have also been used in spaceflight experiments, mice offer closer genetic similarity to humans. This provides more direct insights into human biology than those offered by other model organisms available in space. The rapid development and relatively short lifespan of mice allows observation of space stress-induced adverse effects on shorter time scales than those of humans^18^. The availability of a larger number of mice per mission enhances experimental reproducibility. These advantages have established mice as an essential model animal in space biology research.

To support such studies, the Japan Aerospace Exploration Agency (JAXA) developed a fully equipped experimental platform, the Multiple Artificial Gravity Research System (MARS)^5^. Using the MARS system, as of 2024, nine Mouse Habitat Unit (MHU) missions have been conducted. Findings and achievements of the MHU missions are summarized in Table 1. Throughout these missions, the MARS hardware has been continuously improved, providing valuable insights into biological responses to space stressesors.

**Table 1.**
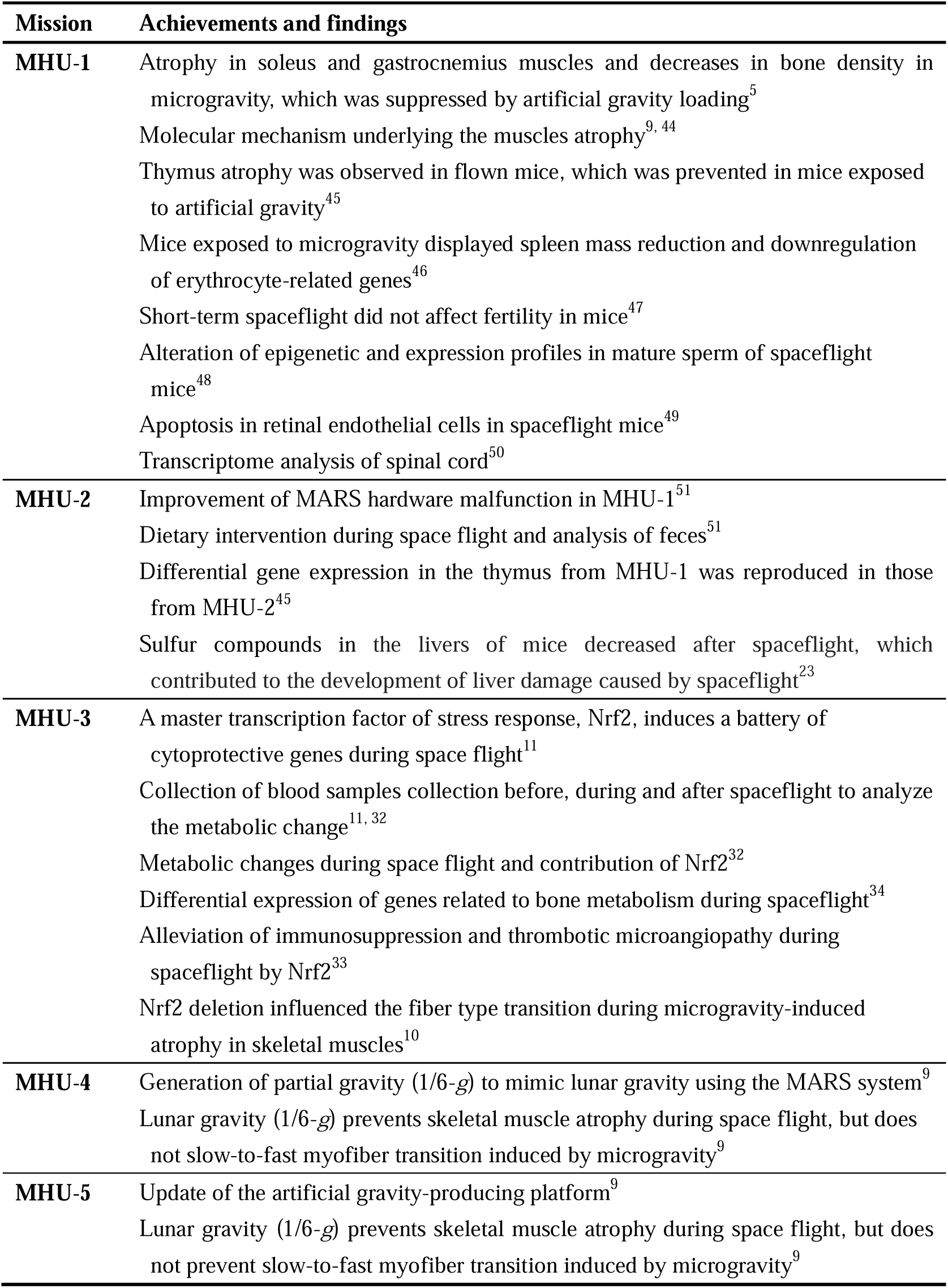
Achievements in MHU missions.

| **Mission** | **Achievements and findings** |
| --- | --- |
| **MHU-1** | Atrophy in soleus and gastrocnemius muscles and decreases in bone density in microgravity, which was suppressed by artificial gravity loading<sup>5</sup><br>Molecular mechanism underlying the muscles atrophy<sup>9, 44</sup><br>Thymus atrophy was observed in flown mice, which was prevented in mice exposed to artificial gravity<sup>45</sup><br>Mice exposed to microgravity displayed spleen mass reduction and downregulation of erythrocyte-related genes<sup>46</sup><br>Short-term spaceflight did not affect fertility in mice<sup>47</sup><br>Alteration of epigenetic and expression profiles in mature sperm of spaceflight mice<sup>48</sup><br>Apoptosis in retinal endothelial cells in spaceflight mice<sup>49</sup><br>Transcriptome analysis of spinal cord<sup>50</sup> |
| **MHU-2** | Improvement of MARS hardware malfunction in MHU-1<sup>51</sup><br>Dietary intervention during space flight and analysis of feces<sup>51</sup><br>Differential gene expression in the thymus from MHU-1 was reproduced in those from MHU-2<sup>45</sup><br>Sulfur compounds in the livers of mice decreased after spaceflight, which contributed to the development of liver damage caused by spaceflight<sup>23</sup> |
| **MHU-3** | A master transcription factor of stress response, Nrf2, induces a battery of cytoprotective genes during space flight<sup>11</sup><br>Collection of blood samples collection before, during and after spaceflight to analyze the metabolic change<sup>11, 32</sup><br>Metabolic changes during space flight and contribution of Nrf2<sup>32</sup><br>Differential expression of genes related to bone metabolism during spaceflight<sup>34</sup><br>Alleviation of immunosuppression and thrombotic microangiopathy during spaceflight by Nrf2<sup>33</sup><br>Nrf2 deletion influenced the fiber type transition during microgravity-induced atrophy in skeletal muscles<sup>10</sup> |
| **MHU-4** | Generation of partial gravity (1/6-g) to mimic lunar gravity using the MARS system<sup>9</sup><br>Lunar gravity (1/6-g) prevents skeletal muscle atrophy during space flight, but does not slow-to-fast myofiber transition induced by microgravity<sup>9</sup> |
| **MHU-5** | Update of the artificial gravity-producing platform<sup>9</sup><br>Lunar gravity (1/6-g) prevents skeletal muscle atrophy during space flight, but does not prevent slow-to-fast myofiber transition induced by microgravity<sup>9</sup> |

**Table 2.** External databases accessible from ibSLS and their target species.

| External databases | Data | Species |
| --- | --- | --- |
| Ensembl | Gene ontology | *Mm, Hs* |
| National Center for Biotechnology Information (NCBI) | Gene ontology | *Mm, Hs* |
| Reference Expresssion Dataset (RefEx) | Gene expression | *Hs* |
| Genotype-Tissue Expression (GTEx) | Gene expression | *Hs* |
| Human Metabolome Database (HMDB) | Metabolite abundance | *Hs* |
| Japanese Multi Omics Reference Panel (jMorp) | Gene expression and Metabolite abundance | *Hs* |

*Mm: Mus musculus; Hs: Homo sapiens*

Mouse experiments enable various large-scale analyses, and the MHU missions have been generating extensive datasets, including transcriptomic, metabolomic, and phenotypic data from spaceflight mice. Moreover, through the space mouse projects, numerous scientific insights have been obtained. However, opportunities to participate in spaceflight projects remain limited because only a small number of experiments can be selected and conducted. Under this situation, we propose establishing a space mouse biobank, which archives experimental data, associated metadata, and unused biological specimens in a standardized and user-friendly format. We believe that such a biobank mice will maximize the scientific accomplishments of MHU missions.

It should be noted that, for many years raw data obtained from each MHU mission have been deposited in public repositories such as the Gene Expression Omnibus (GEO)^2^, the Sequence Read Archive (SRA)^19^ or GeneLab^20, 21, 22^ maintained by the National Center for Biotechnology Information (NCBI) or the National Aeronautics and Space Administration (NASA), respectively. In addition, the biospecimens recovered from the MHU missions have been stored and some made available by JAXA through the MHU Biospecimen Sharing Program (BSP)^23, 24, 25^. However, while these datasets and samples serve as valuable global resources, their broader use across scientific communities remains limited for several reasons. For instance, researchers are needed to equip advanced bioinformatics skills to perform integrative analyses and visualization of raw multi-omics data. Moreover, the limited availability of detailed descriptions of mission design and experimental procedures makes involvement in space life science research challenging. These issues hinder researchers outside the field of space life sciences from fully understanding the dataset context and properly interpreting the results.

To fully realize the scientific potential of the accumulated spaceflight resources, we believe that it is essential to establish a comprehensive platform that integrates datasets, enriches metadata descriptions, enables user-friendly data exploration and analysis, and facilitates the sharing of biospecimens to support further research inspired by past spaceflight missions. To address these challenges, we have developed the integrated biobank of Space Life Science (ibSLS; https://ibsls.megabank.tohoku.ac.jp). The ibSLS aims to enhance the scientific value of the MHU missions and broaden participation in space-based life science research by providing a platform for the utilization of archived datasets and previously unexamined biospecimens.

## Dataset in ibSLS

As of 2025, ibSLS contains transcriptome profiling data from 291 samples based on RNA sequencing (RNA-Seq) analysis derived from MHU-1 through MHU-5 and metabolome profiling data from 228 samples based on either liquid chromatography-mass spectrometry (LC-MS) or nuclear magnetic resonance (NMR) analyses derived from MHU-3 through MHU-5 (Fig. 1a). Transcriptome data cover a wide range of tissues, including nervous tissues (brain and spinal cord), hematopoietic tissues (spleen and bone marrow), immune tissues (thymus and lymph nodes), liver, kidney, brown and white adipose tissues, bone, and muscles (Fig. 1b). The metabolome dataset comprises analyses of plasma samples collected from the inferior vena cava or tail vein, which were collected in MHU-3, MHU-4, and MHU-5 (Fig. 1b). By leveraging the mouse as a model organism, users can utilize data obtained from various tissues to investigate the molecular mechanisms underlying spaceflight-induced phenotypes.

**Figure 1.**
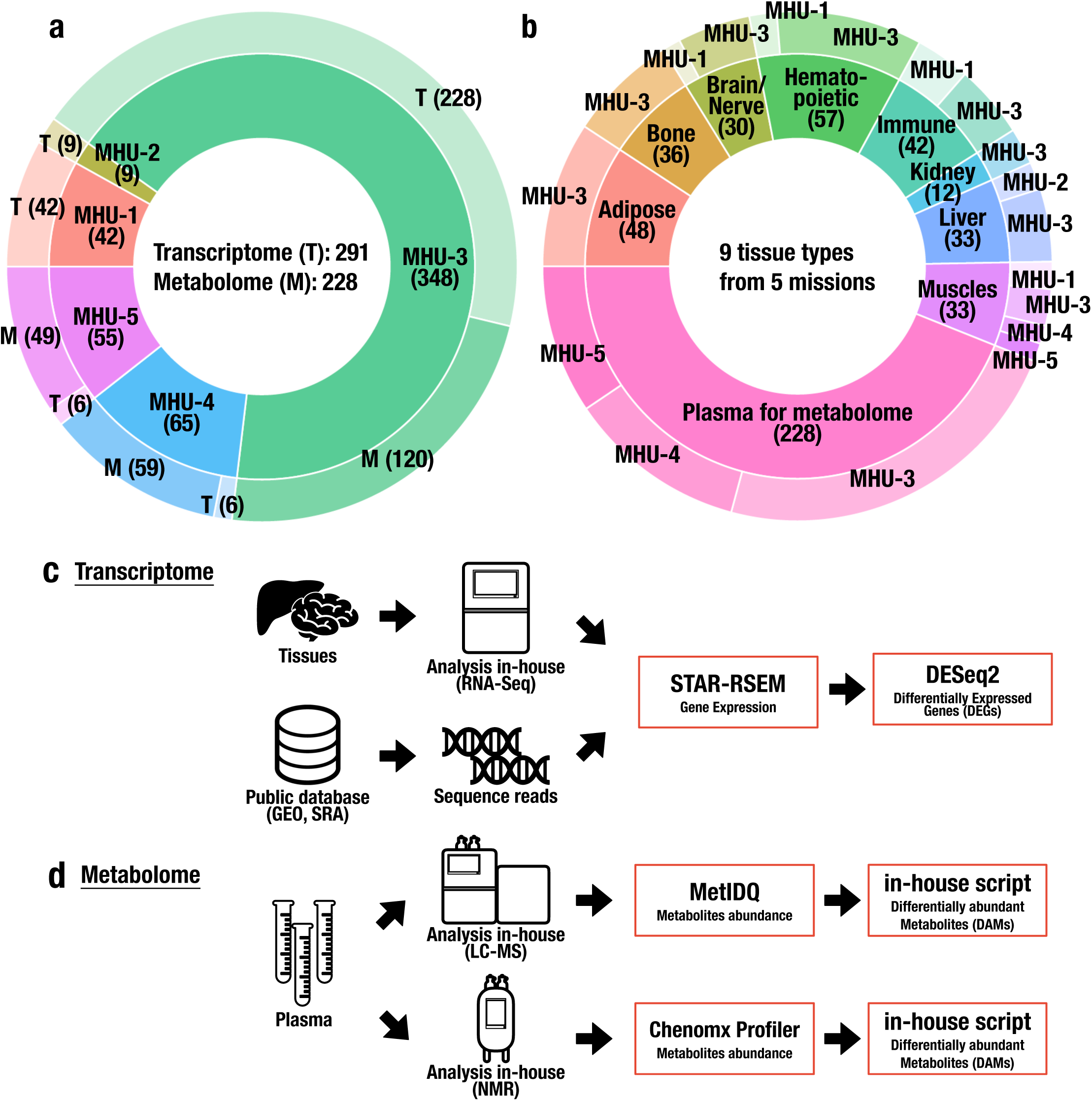
Overview of the ibSLS database content. **a.** Numbers of samples available in the current ibSLS, categorized by assay type (outer ring) and mission (inner ring), with sample counts indicated in parentheses. **b.** Numbers of samples available in the current ibSLS, categorized by mission (outer ring) and tissue type (inner ring), with sample counts indicated in parentheses. **c and d.** Workflow for transcriptome (**c**) and metabolome (**d**) dataset construction. Numerical data visualized in ibSLS are highlighted in red rectangles.

RNA-Seq datasets were constructed from two sources, one from analyses using in-house sequencing facilities and the other from public databases, including GEO and SRA (Supplementary Table S1). To ensure reproducibility and comparability of the data, we used a consensus pipeline, referencing protocols established by groups such as ENCODE^26^ and NASA GeneLab^27^. Raw reads were aligned to the mouse mm10 genome using STAR software^28^ implemented in RSEM software^29^, and transcripts per million (TPM) values were simultaneously obtained for each gene (Fig. 1c). Once the reads were mapped and quantified, differential expression analysis was performed using the DESeq2^30^. The resulting gene-expression tables and lists of differentially expressed genes were deposited and visualized in ibSLS.

To construct the metabolomic dataset, we used two analytical platforms: LC-MS and NMR spectroscopy (Fig. 1d). To ensure comparability across missions and datasets, we generated all metabolomic data in-house using standardized protocols established for a large human cohort project^31^. LC-MS-based profiling was performed using the MxP Quant 500 kit (Biocrates), which enables quantification of 624 metabolites across 26 classes (Supplementary Table S2), allowing comprehensive metabolite profiling and direct mouse-human comparisons. We also conducted NMR-based metabolomic profiling. Although this method requires more plasma than the LC-MS-based method and measures a limited set of 40 metabolites (Supplementary Table S3), it generally provides higher accuracy and reproducibility than the LC-MS-based approach. Differentially abundant metabolites (DAMs) were identified using custom in-house scripts and incorporated into ibSLS.

### Architecture of ibSLS and its function

The overall structure of the ibSLS website is shown in Figure 2a. To design a comprehensive and user-friendly platform, ibSLS incorporates three key functions that facilitate the efficient utilization of spaceflight-related biological data. First, the database provides detailed information about each mission (highlighted in red), including its objectives, experimental design, and key publications. With this function, researchers can fully understand the context in which the data were generated.

**Figure 2.**
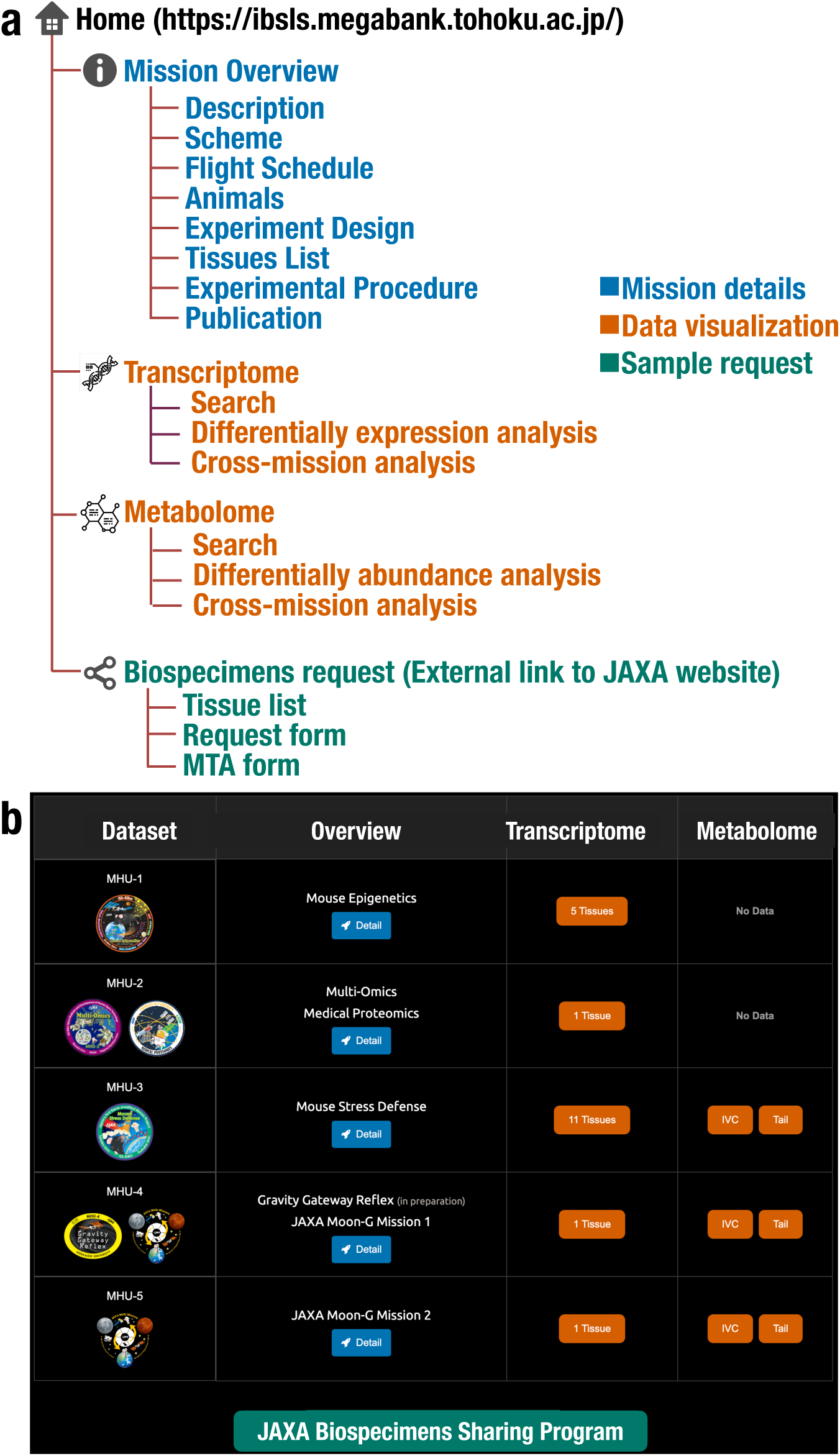
Overview of ibSLS functions. **a.** Structure of the ibSLS website. The web interface is composed of three sections: mission descriptions (blue), data search and visualization (orange), and sample request (green). **b.** Snapshot of the homepage, showing intuitive navigation across all available functions

Second, ibSLS enables visualization of multi-omics data obtained from each mission, allowing intuitive comparisons across missions and conditions (highlighted in orange). To this end, we employed modern web technologies: the Bootstrap CSS framework, whitch ensures a responsive design, and interactive data visualizations and processing are rendered with the Highcharts JavaScript library. Such comparative analysis is crucial for identifying conserved molecular patterns and understanding biological responses to spaceflight. Third, the database provides access to biological samples for their studies (highlighted in green), promoting further investigations and potential collaborations.

All these functions are easily accessible from the ibSLS homepage (Fig. 2b), which does not require registration or authentication to ensure open access to the scientific community. The ibSLS platform is accessible in all major modern web browsers, ensuring a consistent user experience across different systems. We verified compatibility with Google Chrome, Microsoft Edge, Mozilla Firefox, and Apple Safari.

To maintain the integrity and transparency of the datasets, we curated and archived comprehensive information for each dataset. For example, information for MHU-1 is shown in Figure 3. This includes scientific objectives and technical setups of each mission, as well as timelines from launch to return and detailed flight schedules, providing essential temporal context for interpreting experimental outcomes.

**Figure 3.**
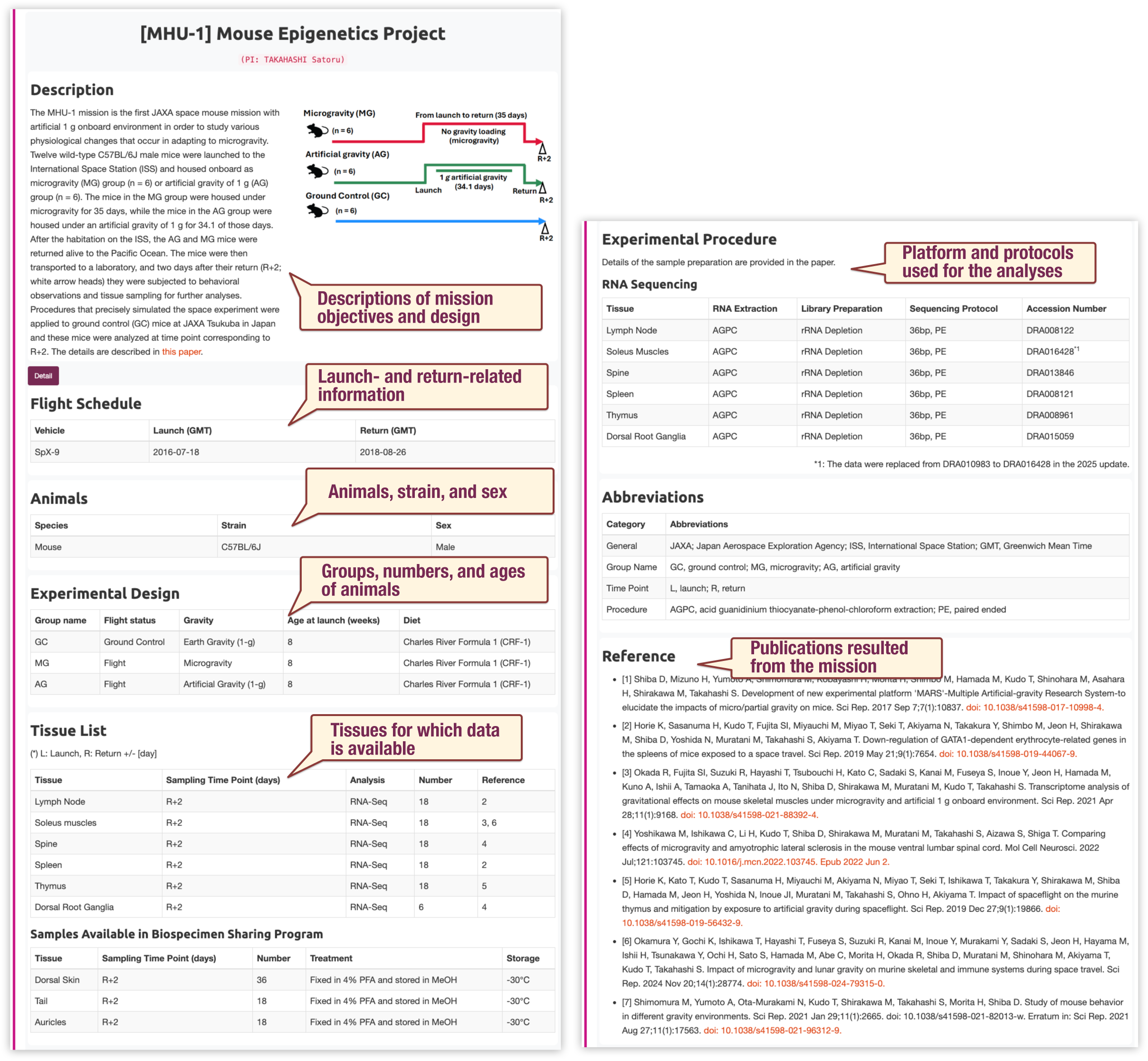
Snapshot of a mission detail page. Example of a detailed mission description. The key experimental conditions and contextual details are highlighted.

Information on the animals and experimental design, such as strain, sex, genotype, age at launch, and grouping based on flight status (*e.g.* flight or ground control) and mission-specific interventions (*e.g.* gravity loading or dietary manipulation), is also included. At the sample level, we documented time points for sample collection and types of analyses conducted on the samples. Protocols for RNA-Seq and metabolome analyses are clearly described with accession numbers for raw data deposited in public repositories to ensure reproducibility. In addition to analyzed data, ibSLS catalogs available biospecimens through the JAXA biorepository, with accompanying details on sample collection and storage conditions. This structured documentation enables users to assess the reliability and applicability of the datasets.

### Visualization of transcriptome and metabolome data in ibSLS

One of the primary and salient functions of ibSLS is the visualization of spaceflight-related multi-omics data. To this end, we implemented a function that allows users to interactively explore transcriptome and metabolome profiles (Fig. 4). By linking these profiles to human multi-omics datasets available from external source, ibSLS facilitates the identification of conserved biological responses, which enables a deeper understanding of spaceflight-related phenotypes and their relevance to human health.

**Figure 4.**
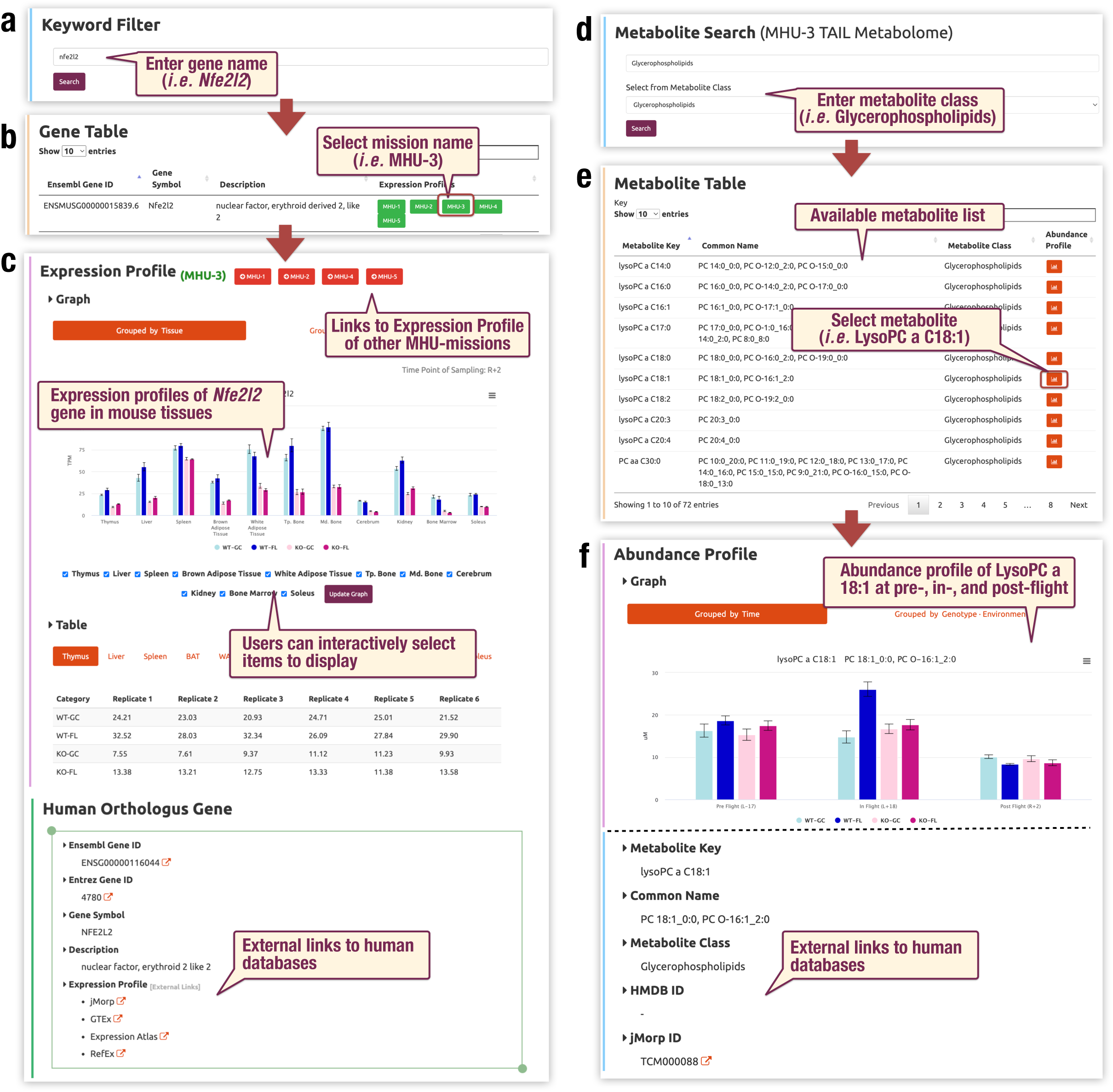
Search function for gene expression and metabolite abundance profiles. **a-c.** Using the *Nfe2l2* gene as an example, this section shows how the results of transcriptome analysis can be visualized. Users can search transcriptome data by entering a mouse gene name or relevant keyword (**a**). Matching results link to MHU mission-specific pages where expression levels of *Nfe2l2* are displayed (**b**). A bar graph shows *Nfe2l2* expression profiles across tissues. Bars represent mean ± s.d. Light blue, dark blue, pink, and magenta bars indicate expression levels in wild-type ground control (GC-WT), wild-type flight (FL-WT), Nrf2-knockout ground control (GC-KO), and Nrf2-knockout flight (FL-KO) mice, respectively. Information on the human ortholog of the mouse gene and external links to human databases are also provided (**c**). **d–f.** Using the metabolite “LysoPC a C18:1,” a member of the glycerophospholipid family, as an example, this section illustrates visualization of metabolome data. Users can search by metabolite name or class (**d, e**). The abundance profile of “LysoPC a C18:1” is shown as a bar graph (**f**). Bars represent mean ± s.d. Bar colors are consistent with those in panel **c**.

To illustrate this function, we employed the case of *Nfe2l2* gene which encodes Nrf2, a master transcription factor involved in oxidative stress response. In the MHU-3 mission, we conducted a one-month spaceflight experiment using both Nrf2-knockout and wildtype mice^11^. Comprehensive analyses of transcriptome and metabolome data demonstrated that Nrf2 plays a central role in adaptive responses to the spaceflight environment, including metabolic adaptation^32^, immune modulation^33^, bone metabolism^34^, and muscle remodeling^10^ under microgravity conditions.

Using this case as an example, Figure 4 illustrates how users can visualize gene expression data. Genes of interest can be searched by gene names or related keywords (Fig. 4a). When matching entries are found, links are provided to gene-specific pages across each MHU mission (Fig. 4b), where expression profiles across tissues analyzed in the mission are displayed as bar graphs (Fig. 4c). These pages feature interactive controls for selecting tissues and conditions, allowing users to generate customized visualizations suited to their research interests. To facilitate cross-species comparisons and integrate the mouse dataset with large-scale human datasets, we annotated mouse genes with their corresponding human orthologs. We established links to external resources containing extensive gene expression profiles, as well as large cohort-based human phenotype data (Fig. 4c). These human datasets include the Reference Expression Dataset (RefEx)^35^ for standardized gene expression across tissues, the Genotype-Tissue Expression (GTEx)^36^ for tissue-specific gene expression variability, and the Japanese Multi-Omics Reference Panel (jMorp) described below.

A similar framework is provided for metabolome data visualization. We implemented a function that allows users to select queries from a list of available metabolites or metabolite classes (Fig. 4d and 4e). Since metabolite nomenclature is not always standardized, this feature ensures easier access to the desired metabolite. As an example, we illustrated how to search metabolomic profiles (Fig. 4d and 4e). Users can first select a metabolite class such as *glycerophospholipids*, and then choose a specific metabolite of interest, *LysoPC a C18:1* in this case, to view its abundance profile. The abundance profiles of metabolites in multiple time points are presented as bar graphs for clear visualization (Fig. 4f). The mouse metabolites were annotated to human metabolites using the Human Metabolome Database (HMDB)^37^.

### Link to human data in jMorp

To enhance translational research across species, we linked transcriptome and metabolome data in ibSLS to external human databases, including the Japanese Multi-Omics Reference Panel (jMorp) (Table2). jMorp is a database that we have been constructing to provide integrated genomic, transcriptomic, and metabolomic data from large-scale human cohorts in the Tohoku Medical Megabank Project in Japan^12, 13, 16^. jMorp comprises multi-omics profiles from healthy volunteers across a broad age range, accompanied by extensive phenotype information, including physiological, biochemical, and lifestyle-related parameters. By linking mouse datasets with real-world human datasets within a unified platform, users can evaluate whether spaceflight-induced molecular changes in mice correspond to variation across age or phenotype in humans.

### Function for statistical analysis of transcriptome and metabolome data

ibSLS also offers several advanced analytical functions to facilitate in-depth exploration of spaceflight-induced molecular changes. One of its key features is the ability to identify differentially expressed genes (DEGs) and differentially abundant metabolites (DAMs), which are critical for understanding biological responses to the space environment. This function allows researchers to systematically analyze how different spaceflight conditions affect gene expression and metabolic profiles.

As an example of usage, Figure 5 shows how to visualize gene expression changes that occurred in kidney, based on observations in MHU-3 mission^34^. Users can specify a mission (*e.g.,* MHU-3), genotype of mice (*e.g.* wildtype), tissues of interest (*e.g.* kidney) and groups for comparison (*e.g.,* Ground Control [GC] vs. Flight [FL]), and define a custom false discovery rate (FDR) threshold (*e.g.,* FDR = 0.01), allowing flexible identification of DEGs associated with different biological perturbations (Fig. 5a). Based on these parameters, ibSLS generates a comprehensive list of significantly up or downregulated genes (Fig. 5b). Each output includes logLJ-transformed fold change, FDR, and a direct link to detailed gene information.

**Figure 5.**
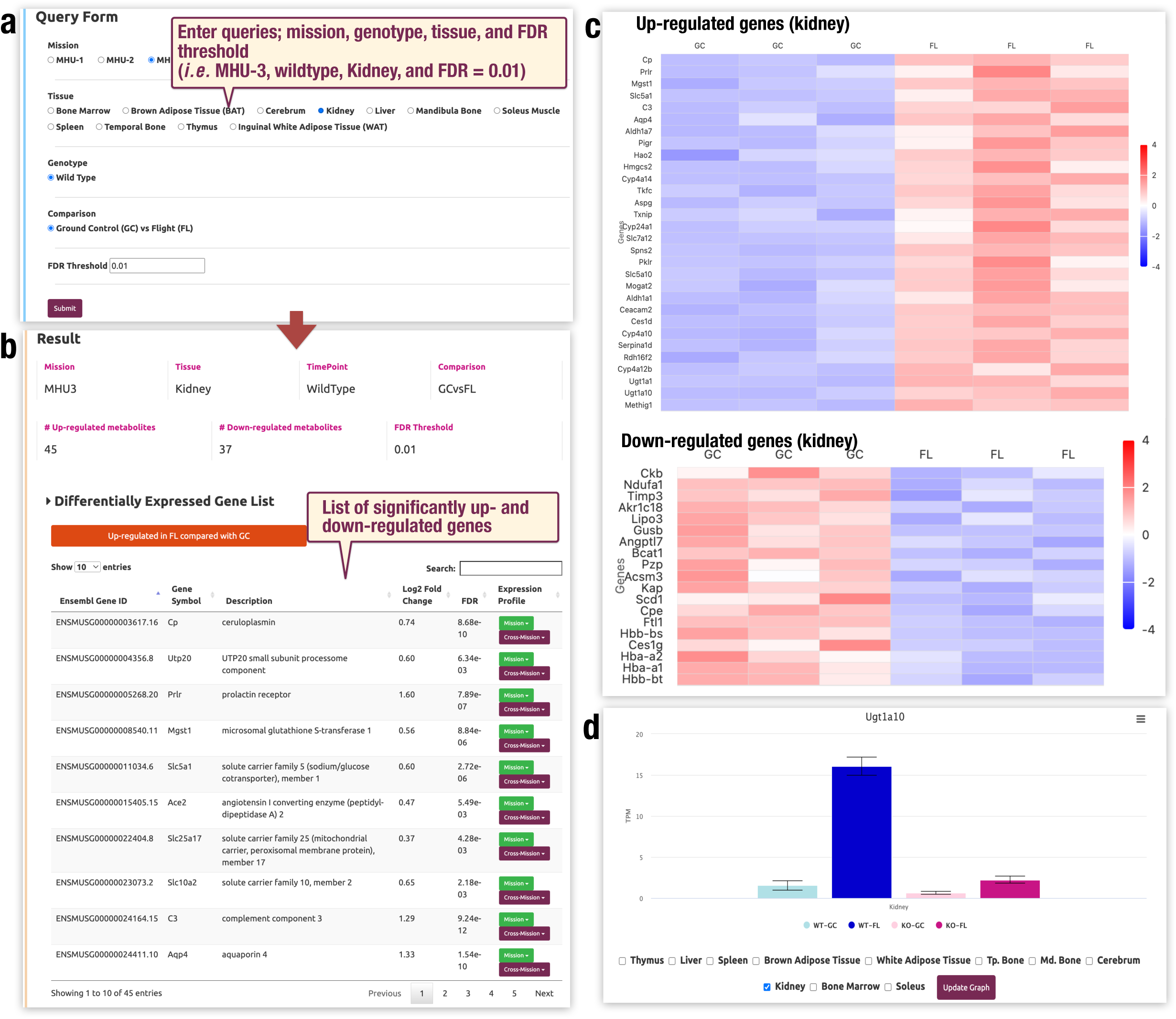
Function to identify differentially expressed genes (DEGs). **a.** Queries required for differential expression analysis. Users select a mission, tissue, group, and custom FDR threshold. **b.** DEG list resulting from the search, showing significantly upregulated and downregulated genes in spaceflight vs. ground control mice. **c.** Heatmap showing expression patterns of DEGs across experimental groups. TPM values were normalized to Z-score for visualization. **d.** Example bar graph showing expression levels of *Ugt1a10* in kidney, which was significantly upregulated in spaceflight mice compared with ground controls and whose upregulation markedly abrogated by Nrf2 knockout. Bars represent mean ± s.d. Light blue, dark blue, pink, and magenta bars indicate expression levels in wild-type ground control (GC-WT), wild-type flight (FL-WT), Nrf2-knockout ground control (GC-KO), and Nrf2-knockout flight (FL-KO) mice, respectively.

To enhance interpretability, the results can be exported to a text file and visualized as a heatmap (Fig. 5c), allowing users to identify patterns and trends across conditions. By clicking an individual gene name, expression profile of the gene appears as a bar graph (Fig. 5d). Notably, this visualization clearly illustrated the attenuation of spaceflight-induced *Ugt1a10* expression in NRF2 knockout mice, consistent with a previous report^34^. In addition, the identified genes are linked to their respective expression observed in different tissues (*e.g.,* expression profile in liver shown in Supplementary Fig. S1), providing seamless access to comparative data. The same function is implemented for metabolome data, which enables users to explore DAM in plasma samples.

### Function for data comparison between missions

We further implemented a function for comparing statistical analysis results across two different missions, termed a “cross-mission comparison analysis” (Fig. 6). This feature facilitates straightforward evaluation of reproducibility between missions and contributes to more reliable data interpretation. The cross-mission comparison analysis requires two sets of input queries, missions, tissues, experimental groups, and a common FDR threshold for both conditions (Fig. 6a).

**Figure 6.**
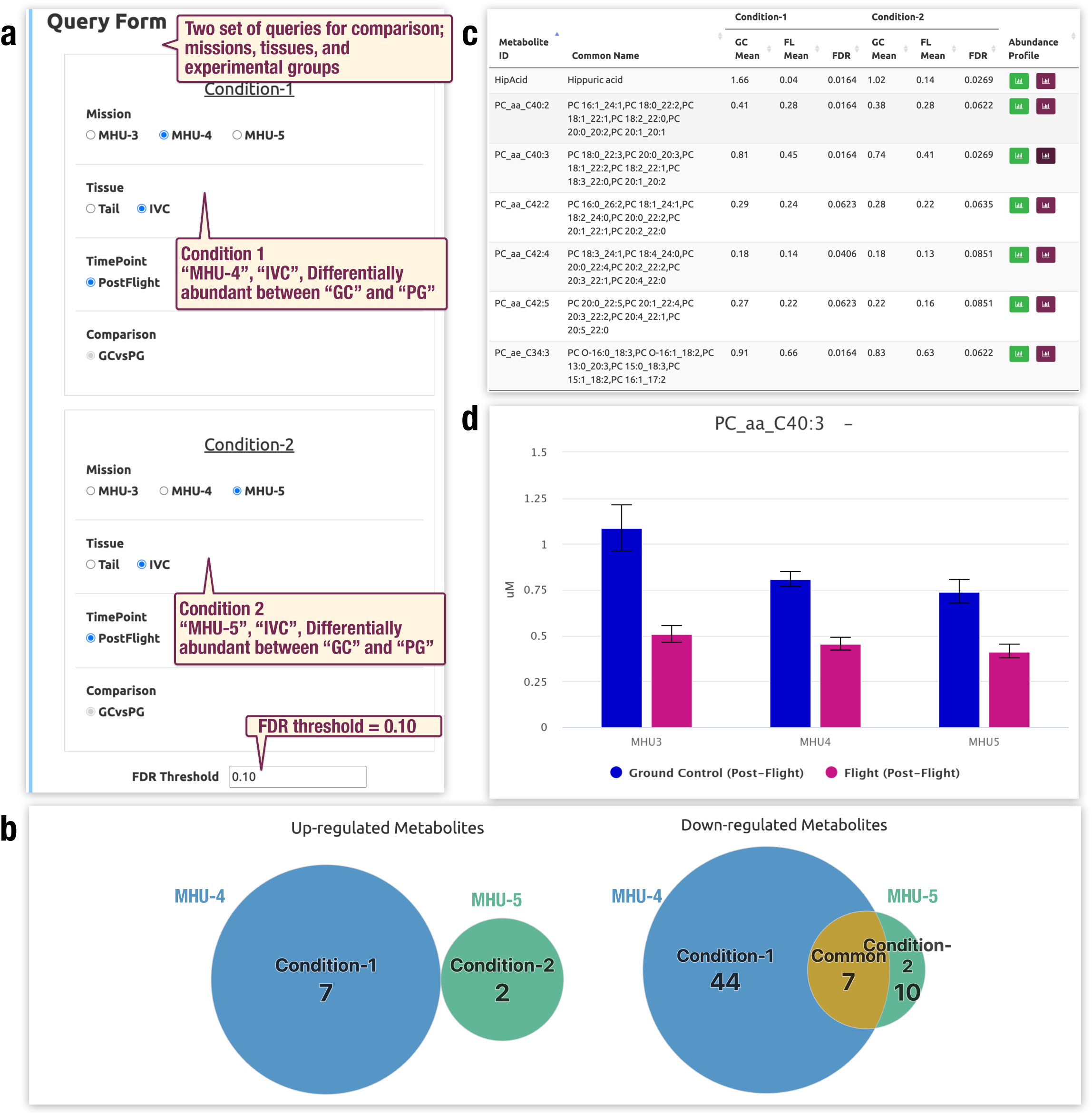
Function to explore reproducibility of findings by comparing results from two missions. **a.** Users can specify two conditions to evaluate differential expression. **b.** Overlaps of upregulated (left) and downregulated (right) metabolites between condition 1 (MHU-4) and condition 2 (MHU-5) are shown as Venn diagrams. **c**. Lists of metabolites specific to each condition and those common to both conditions are generated. Here, the list of metabolites commonly downregulated in MHU-4 and MHU-5 is shown. **d.** Bar graph showing “PC aa C40:3” level in plasma recovered from spaceflight and ground control mice in MHU-3, MHU-4, and MHU-5. Bars represent mean ± s.d. Magenta and blue bars indicate abundance levels in the flight (FL) group and in the ground control (GC) group, respectively.

To demonstrate this function, we assessed the reproducibility of spaceflight-induced metabolic alterations by comparing results from the MHU-4 and MHU-5 missions. Conducted in 2019 and 2020, respectively, both missions followed comparable experimental designs in which mice were maintained under 1/6 g partial gravity (PG) for 30 or 31 days during spaceflight, respectively^9^. For the analysis, we defined a comparison between the metabolic profiles of spaceflight mice under PG and ground control mice (GC) from the MHU-4 mission as “Condition 1” and an identical comparison from the MHU-5 mission as “Condition 2” (Fig. 6a).

Based on these conditions, ibSLS outputs a Venn diagram illustrating the overlap of differentially abundant metabolites (DAMs) between the two conditions (Fig. 6b). In the MHU-4 and MHU-5 missions, seven and two metabolites were upregulated, while 44 and 10 metabolites were downregulated, respectively (Fig. 6b and c). Among these, seven metabolites were reproducibly downregulated in both missions, the majority of which belonged to the phosphatidylcholine (PC), a subgroup of glycerophospholipid (GP) class. Notably, we previously reported downregulation of many GPs in spaceflight mice compared to ground controls in the MHU-3 mission^32^.

Using the cross-mission function in ibSLS, we further assessed whether these GP-related findings were consistently observed in MHU-3 as well (Fig. 6d and Supplementary Fig. S2). This analysis highlights the utility of the ibSLS database in enabling user-friendly, comprehensive cross-mission comparisons. Overall, this integration empowers users to explore MHU datasets holistically, facilitating deeper insights into spaceflight-induced molecular changes.

### Distribution of biospecimens through the MHU Biospecimen Sharing Program

As part of the broader ibSLS initiative, the MHU Biospecimen Sharing Program facilitates broad access to mouse tissue samples collected during past MHU missions. Unanalyzed tissue samples are stored at the JAXA Biorepository (https://humans-in-space.jaxa.jp/kibouser/provide/mhu/71891.html) at JAXA Tsukuba Space Center in Japan. These tissue samples have been provided to investigators in Japan and the United States under the Japan-US Open Platform Partnership Program (JP-US OP3), offering researchers the opportunity to analyze post-flight biological materials. Notably, ibSLS also provides the spaceflight-related biological data for researchers worldwide, enabling new experimental endeavors based on unique ideas.

Through the MHU Biospecimen Sharing Program, researchers can request archived mouse tissue samples remaining from the original MHU missions, eliminating the need for new spaceflight experiments. As of June 2025, the MHU Biospecimen Sharing Program, including the ibSLS tissue sample sharing program, has delivered 134 types of mouse tissue samples for 52 biological studies. These samples were derived from five MHU missions and have supported space biology research on a global scale. Since available tissues, organs, and experimental conditions vary by mission, each sample is accompanied by associated metadata and omics datasets provided through ibSLS. This ensures seamless integration of physical sample access with available digital data.

Overall, ibSLS establishes a comprehensive platform for space life science by integrating detailed metadata, multi-omics datasets, and biospecimen sharing from multiple MHU missions. By democratizing access to both digital resources and physical samples, ibSLS maximizes the scientific value of past spaceflight experiments, fosters international collaboration, and accelerates discoveries on the biological impact of space. This open framework not only supports the development of strategies for sustaining long-duration human space exploration but also advances biomedical knowledge relevant to human health on Earth.

## Discussion

Since its launch in 2020, ibSLS has been providing multi-omics data from MHU missions. In the current update, we have implemented further improvements in data search and integrative analysis functions, as well as seamless access to the sample-sharing platform. By providing a unified platform that combines rich multi-omics data with human phenotype data, ibSLS enables researchers to gain deeper insights from their analyses. While the current ibSLS includes data from MHU-1, 2, 3, 4, and 5 missions, we are planning to expand the database to incorporate data from additional missions (Supplementary Fig. S3). As new datasets become available, we will systematically curate and integrate them to provide a more comprehensive resource for space life science research.

Several platforms have been developed in the global context of data visualization in space life science, including the Space Omics and Medical Atlas (SOMA)^38^ and GeneLab^20, 21, 22^. SOMA primarily focuses on human datasets from missions such as the NASA Twins Study^39^, JAXA’s Cell-Free Epigenome Study^40^, and SpaceX’s Inspiration4^41^. SOMA has also begun to include mouse datasets from NASA GeneLab. A repository of human specimens has been organized by the Cornell Aerospace Medicine Biobank (CAMbank). Similarly, the GeneLab operates a repository that provides space-related data from a wide range of model organisms, including mice, plants, fruit flies, cultured cells, nematodes, and microorganisms. These platforms share the goal of advancing space biology through open data.

We have designed ibSLS to offer additional advantages as a biobank to these platforms. ibSLS provides enhanced interactive tools that allow users to intuitively explore transcriptomic and metabolomic datasets across multiple missions. In addition, by linking to the large human cohort database developed by the Tohoku Medical Megabank Project, ibSLS supports the interpretation of spaceflight-induced molecular changes of mice in the context of human phenotypic variation, facilitating translational insights from animal models to human health.

Due to the operational limitations inherent in individual spaceflight mouse missions, it is often challenging to fully standardize sample collection as well as analytical procedures across missions. Therefore, we emphasized the importance of descriptions on background information, in addition to the elaborate visualization of multi-omics data. To facilitate accurate interpretations when comparing datasets from different missions, ibSLS provides detailed contextual information associated with each dataset and biospecimen, which enables users to construct hypotheses based on the data available in ibSLS with careful consideration of the background context. We believe that the well-organized information serves as a valuable resource for researchers new to the field of space life science.

We aim to expand ibSLS continuously both by adding data from new MHU missions and by incorporating data from advanced analyses. In case of the latter, future updates will include epigenetic modification data as well as detailed mouse phenotype information. By integrating these data, ibSLS will provide a more comprehensive framework for understanding how molecular changes translate to organism-level responses in spaceflight. To support these datasets, we are also planning to enhance visualization tools to allow users to intuitively explore region-specific chromatin modifications. These expansions will significantly improve the interpretability of omics findings and facilitate multi-layered investigation of biological adaptation to the space environment.

In conclusion, ibSLS plays a pivotal role in advancing space biology by enabling integrative analysis of spaceflight-derived mouse omics data and access to associated biospecimens. Through ongoing expansion of its datasets, we believe that ibSLS will contribute to the democratization of space life science by making high-quality data more accessible and reusable to broader scientific communities.

## Methods

### RNA-Sequencing data

FASTQ files were downloaded from SRA (Accession numbers; DRA008122, DRA010983, DRA013846, DRA008121, DRA008961, DRA011091, and DRA014378) and GEO (Accession numbers; GSE240654 and GSE152382). Raw reads were mapped to the mouse mm10 genome using STAR^28^ (version 2.6.1). Transcripts per million (TPM) values were calculated to measure gene expression using RSEM^29^ (version 1.3.1). STAR mapping was executed through the RSEM toolkit using default parameters. Differentially expressed genes (DEGs) were identified using DESeq2 software^30^.

### Metabolome analysis

Plasma samples from tail blood (5 μL per analysis) were analyzed using the MxP^®^Quant 500 Kit (Biocrates Life Sciences) and the ultra-high-performance liquid chromatography (UHPLC)-MS/MS analyses (Xevo TQ-S system, Waters). For NMR-based metabolome analysis, 50 µL of plasma per sample were subjected to a standard methanol extraction procedure. NMR experiments were performed at 298 K on a Bruker Avance 600-MHz spectrometer equipped with a CryoProbe and a SampleJet sample changer. All data were processed using the Chenomx NMR Suite processor module (Chenomx). Metabolites were identified and quantified using the target profiling approach implemented in the Chenomx Profiler module.

### Database implementation and data availability

The ibSLS web platform was developed using the Django web framework^42^. Data storage is managed with an embedded SQLite database^43^ for simplicity and compatibility. The Bootstrap CSS framework was used fo implement a responsive client-side interface, and interactive data visualizations were rendered using the Highcharts JavaScript library. All figures and numerical data generated by ibSLS are freely available to users, provided that the source is appropriately cited. Detailed guidelines for citation and data usage are outlined in the Terms of Use on our website (https://ibsls.megabank.tohoku.ac.jp/condition_of_use). To monitor usage and improve the service, anonymized access logs are recorded via Google Analytics, which tracks user interactions.

## Acknowledgments

We thank Mr. Sho Furuhashi and Mr. Tetsuo Chiba for technical assistance, and Ms. Yoko Kitami of JAXA for editorial support. This work was supported by the JSPS Grant-in-Aid for Transformative Research Areas (Grant Number 25H013740) and a commissioned research fund from JAXA. It was also partly supported by JSPS KAKENHI (Grant Number 25H01374 to A.O. and 25H01370 to T.S.). All computational resources were provided by the Tohoku University Tohoku Medical Megabank Organization supercomputer system, which is supported by the infrastructure development for large-scale analysis utilizing Genome, and Health and Clinical Data (Grant Number JP21tm0424601).

## Author Contribution

A.O., Y.A., R.O., D.S., and M.Y. wrote the manuscript.

Y.A., A.O., and A.U. designed and developed the user interface of ibSLS. A.O., T.S., and F.K. conducted the transcriptome analysis.

E.H., A.U. and S.K. conducted the metabolome analysis.

S.K., F.K., and K.K. supervised the metabolome, transcriptome, and bioinformatics analyses, respectively.

M.Y. supervised the overall project.

All authors discussed the results and commented on the manuscript.

## Legends to Supplementary Figures

**Supplementary Figure S1.**
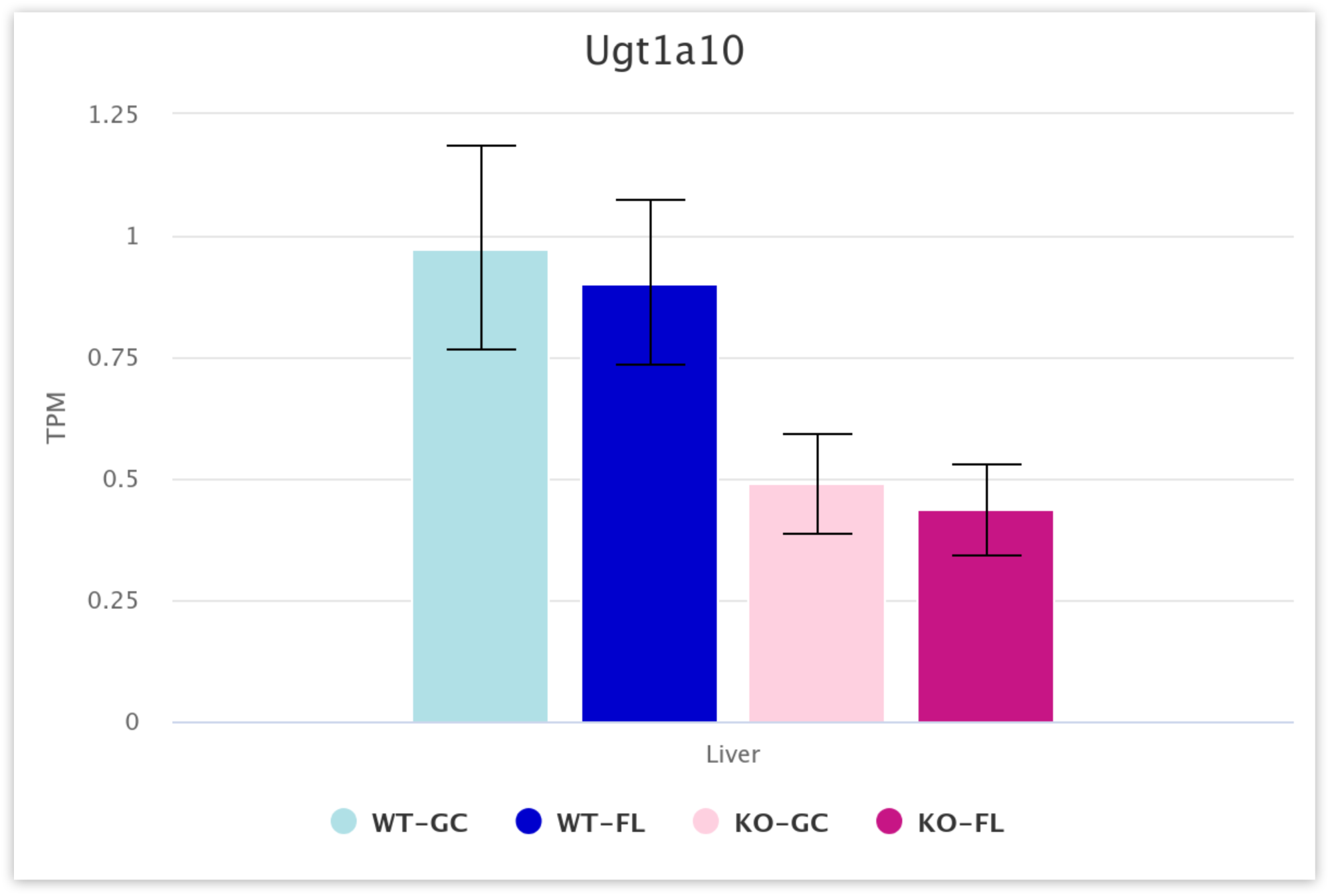
(related to Figure 5d). Bar graph of *Ugt1a10* expression in liver. This view can be directly accessed from the kidney expression analysis page shown in Fig. 5d. Bars represent mean ± s.d. Light blue, dark blue, pink, and magenta bars indicate expression levels in wild-type ground control (GC-WT), wild-type flight (FL-WT), Nrf2-knockout ground control (GC-KO), and Nrf2-knockout flight (FL-KO) mice, respectively.

**Supplementary Figure S2.**
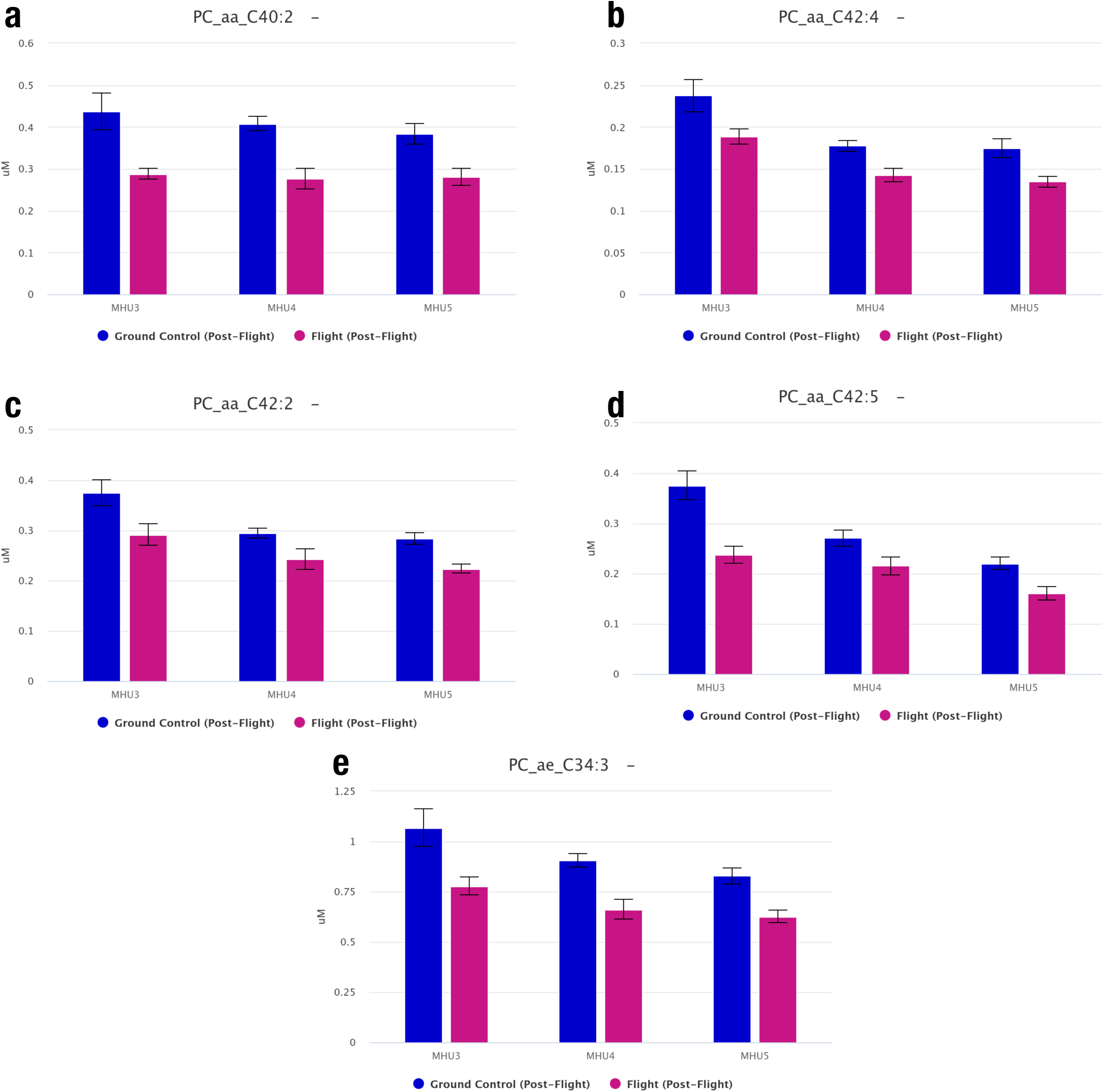
(related to Figure 6d). Downregulation of metabolites belonging to the glycerophospholipid class. Abundance levels of “PC aa C40:2” (**a**), “PC aa C42:4” (**b**), “PC aa C42:2” (**c**), “PC aa C42:5” (**d**), and “PC aa C34:3” (**e**) are shown. Bars represent mean ± s.d. The blue and magenta bars indicate abundance levels in ground control (GC) and flight (FL) group, respectively.

**Supplementary Figure S3.**
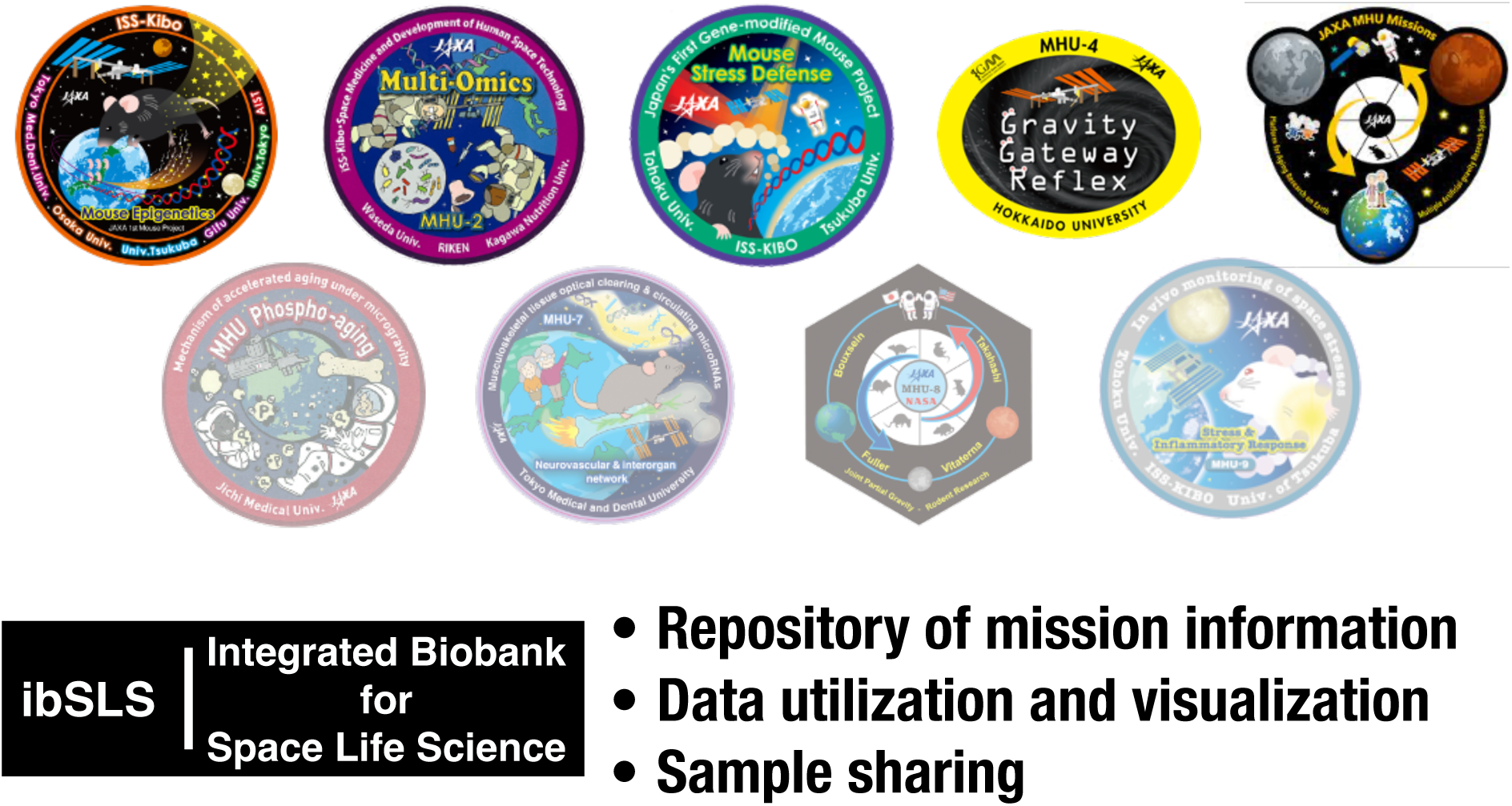
Key concepts of ibSLS. The ibSLS platform integrates detailed metadata, multi-omics data visualization, and biological sample distribution for the MHU spaceflight missions.

